# Effect of sampling media and culture conditions on dendritic cell generation: ion depleted plasma negatively influences the survival of monocyte-derived dendritic cells

**DOI:** 10.1101/066233

**Authors:** Z. Parackova, J. Kayserova, A. Sediva

**Author notes:** Email addresses of authors: Zuzana Parackova Jana Kayserova Anna Sediva. Corresponding author: Parackova Zuzana Department of Immunology, Charles University, 2^nd^ Faculty of Medicine, University Hospital Motol, Prague, V Uvalu 84 150 06 Prague Czech Republic.

## Abstract

**Abbreviations:**

EDTA: ethylenediaminetetraacetic acid
FBS: fetal bovine serum
MFI: mean fluorescence intensity
moDC: monocyte-derived dendritic cells
PBMC: peripheral blood mononuclear cells

Autologous serum or plasma are often used as a supplementation in culture media for autoimmune disease studies. They contain many biologically and immunologically active molecules that might be involved in the development of these diseases. We examined the effect of different cultivating media supplemented with autologous serum and two types of plasma obtained from heparin and EDTA coated tubes on monocyte-derived dendritic cells viability. Our results show that medium with plasma from EDTA coated tubes is unsuitable for cultivation of myeloid cells due to the absence of bivalent ions and that the type of sampling tubes used for blood collection might be critical for successful experiment.

## Introduction

Autologous serum or plasma are commonly used as a substitution for the universally used fetal bovine serum (FBS) in experimental applications for different types of cells. FBS is not an ideal form of supplementation because the full composition of FBS is not known and its component concentrations can change from batch to batch. FBS may contain pathogenic viruses or prions. Additionally, immunological complications may occur since FBS contains proteins that induce immune reactions^1^. Instead, autologous serum or plasma are commonly used in the generation of monocyte-derived dendritic cells (moDC), especially in clinical application studies^2–7^. Autologous plasma or serum, due to their content of many important disease-modifying factors, also form a crucial part in the experimental setting when cell culture is used in order to study disease mechanisms^8–11^. Our aim was to study the effect of autologous plasma or serum on moDC functional properties. During our initial experiments we noticed that moDC generated in autologous plasma had markedly decreased viability. Thus, we tested different culture conditions in order to optimize the moDC cultivation and obtain optimal and trustworthy experimental results.

## Material and Methods

### Plasma, serum and media preparation

Plasma and serum were obtained from peripheral blood of healthy volunteers. Serum was collected from coagulated blood (Greiner Bio-One, Kremsmünster, Austria). Plasma was obtained from uncoagulated blood EDTA-coated tubes (Greiner Bio-One) (Scheme 1). The tubes were centrifuged 3000rpm, 10min, 25°C. Plasma and serum were aliquoted and stored in −80°C. For experiments complete media was used containg RPMI 1640 (ThermoFischer Scientific, Waltham, MA, USA), penicillin and streptomycin, glutamin (ThermoFischer Scientific) and supplemented either with autologous serum or plasma or fetal bovine serum (GE Healthcare Life Sciences).

**Scheme 1:**
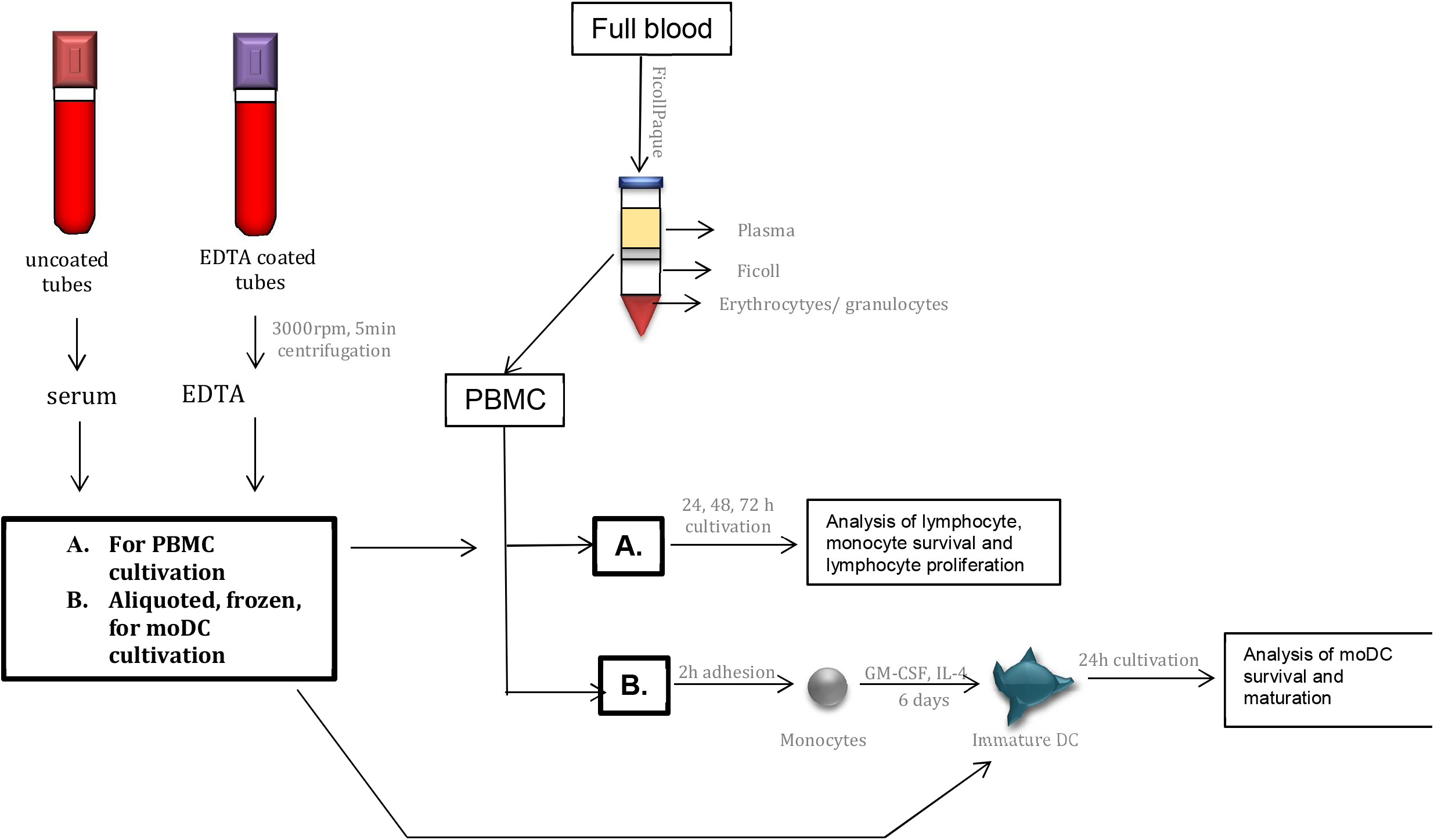
Autologous plasma and serum were obtained from peripheral blood of healthy volunteers by centrifugation. 10% serum or plasma supplementation was used for (A.) PBMC cultivation and (B.) monocyte-derived dendritic cells cultivation for 24 hours.

### Generation of monocyte-derived dendritic cells (moDC)

PBMCs (peripheral blood mononuclear cells) were isolated from the peripheral blood of healthy volunteers using the Ficoll-Paque gradient (GE Healthcare Life Sciences AB, Uppsala, Sweden) method. Monocytes were separated by 2h adhesion in culture flasks (Thermo Scientific, Roskilde, Denmark) in density 10^6^cells/cm^2^. Cells were cultivated in complete media supplemented with 10% FBS. moDCs were generated from monocytes for 6 days in complete media in the presence of 40ng/ml IL-4 (CellGenix) and 500IU/ml GM-CSF (CellGenix). Medium and cytokines were replenished on day 3. On day 6, immature dendritic cells were harvested and counted. Immature moDC (iDC) were seeded into 96-wells (Thermo Scientific) in concentration 10^6^/ml in RPMI media without any serum or plasma supplementation. Then, 10% autologous plasma, serum or FBS were added to culture media (Scheme 1). For moDC maturation, iDCs were stimulated with 100ng/ml LPS (Sigma Aldrich, St. Louis, Missouri, USA) for 24hours. When indicated, to conditions where autologous plasma from EDTA-coated tubes was used, ions were added, namely 0.9mmol/l Mg (HBM Pharma, Martin, Slovakia) and 2mmol/l Ca (B Brown, Melsungen, Germany). After 24 hours, moDC were stained with antiCD11c APC (Exbio, Prague, Czech Republic), CD14 PEDy590 (Exbio), CD80 FITC (Beckman Coulter, Marseille, France), CD86 PECY5 (BD Bioscience, San Jose, CA, USA), HLA-DR PECY7 (BD Bioscience) and DAPI (ThermoFisher Scientific). Viability of moDC was analysed as DAPI negative of CD11c positive moDC. The samples were collected using a BD FACSAria II (BD Biosciences,), and BD FACSDiva™ software (BD Biosciences) was used for signal acquisition.

### Cultivation of PBMC

PBMCs (peripheral blood mononuclear cells) were isolated from the peripheral blood of healthy volunteers using the Ficoll-Paque method and seeded in 10% FBS, autologous plasma or serum supplemented RPMI media in concentration 10^6^cells/ml (Scheme 1). When indicated, to conditions where plasma from EDTA-coated tubes were used, 0.9mmol/l Mg; 2mmol/l Ca were added. Percentage of lymphocytes and monocytes of total events was analysed by flow cytometry after 24 and 48 hours.

Furthermore, after isolation lymphocytes were seeded in RPMI supplemented with autologous serum, plasma or FBS and stimulated with antiCD3/CD28 (Exbio) antibodies for proliferation assay. After 3 days, lymphocytes were stained with antiCD4 PB (Exbio), anti CD8 FITC (Exbio) and then were processed using the eBioscience Fixation/Permeabilization solution (eBioscience, San Diego, CA, USA) and intracellular stained for Ki67 PE (BioLegend, San Diego, CA, USA).

## Results and Discussion

To determine the suitable conditions for moDC cultivation and maturation, we decided to study the effect of a supplementation of human autologous serum and plasma to conditioning media (Scheme 1). We collected plasma from EDTA coated tubes, as well as serum from centrifuged coagulated blood. After 6 days of moDC generation in complete media supplemented with FBS, we cultivated moDC in RPMI medium supplemented with 10% autologous serum, plasma or FBS for 24 hours. We found out that moDC viability was higher when culture media was supplemented with FBS or autologous serum and significantly lower when autologous plasma from EDTA coated tubes was used (Figure 1A).

**Figure 1:**
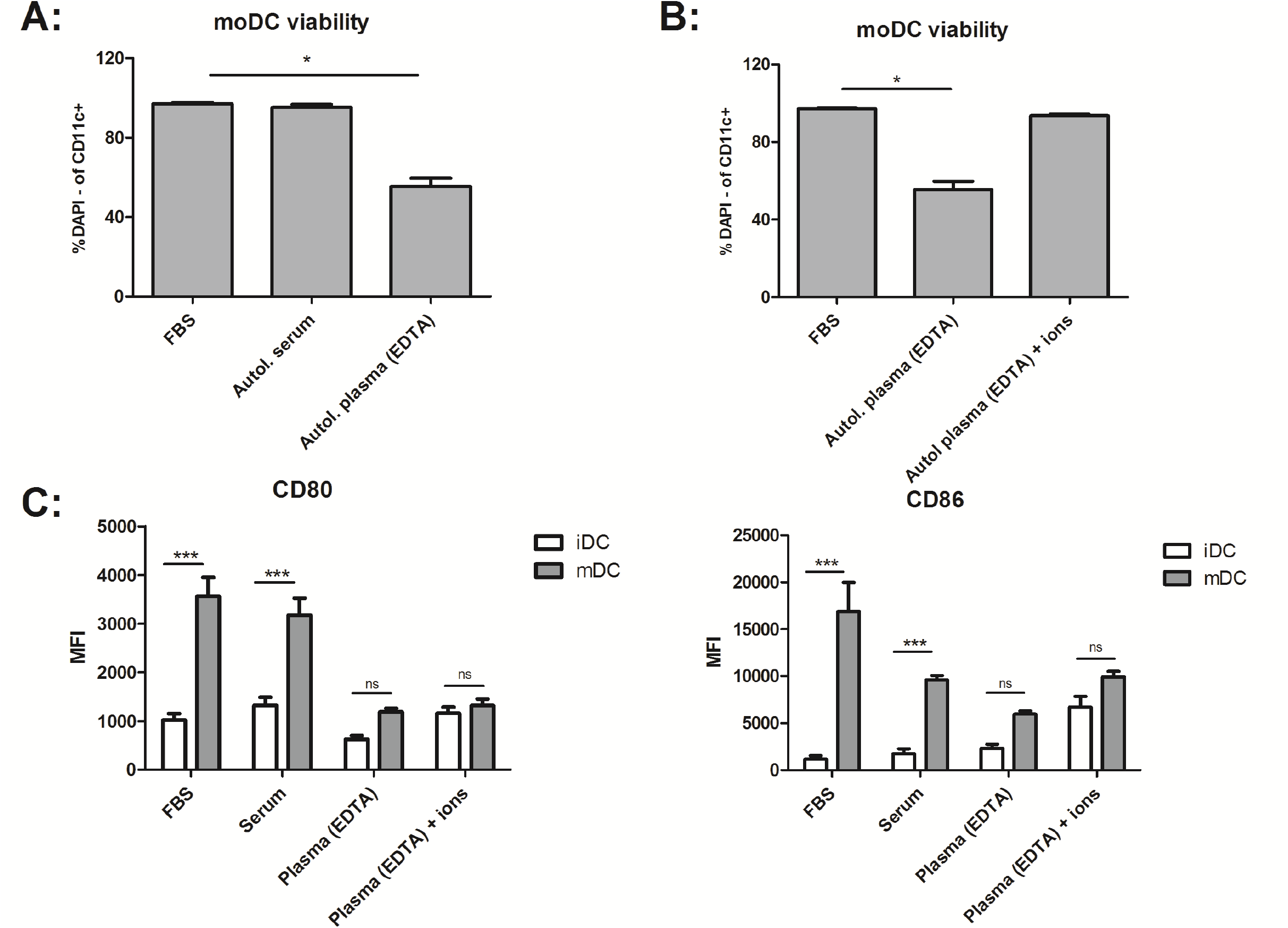
(A) Immature monocyte-derived dendritic cells (moDC) were cultivated in RPMI media supllemented with FBS, autologous serum or plasma and moDC viability was analyzed after 24 hours. Live cells were considered as DAPI negative CDllc possitive moDCs. (B) moDCs viability was tested also after bivalent ions - Ca and Mg were added to the ion-depleted plasma from EDTAcoated tubes. (C) Upregulation of maturation markers CD80 and CD86 (expressed as MFI-mean flourescence intensity) was analyzed after 24 hours of LPS stimulation. The assay was performed in quadruplicates and mean values+/−SEM are shown.

EDTA is a common anticoagulant and chelating agent that binds bivalent ions and therefore plasma from EDTA coated tubes lacks Mg^2+^ and Ca^2+^. The serum and plasma were then analyzed for exact concentration of several ions to establish differences between them. As expected, plasma from EDTA-coated tubes contained almost zero levels of Mg^2+^ and Ca^2+^. We, subsequently, tested the effect of Mg^2+^ and Ca^2+^ by adding them to the plasma obtained from EDTA coated tubes, which was most ion-depleted. When moDC were cultured in media containing 10% plasma supplemented with ions, they retained viability comparable to those cultivated in complete media (Figure 1B).

We also tested the effect of autologous serum and plasma on moDC maturation after lipopolysaccharide stimulation, expressed as MFI (mean fluorescence intensity) of CD80 and CD86 markers. moDCs were not able to upregulate their maturation markers when the ion-depleted plasma was used. Maturation capacity was not restored when ions were added to the plasma supplemented medium (Figure 1C).

Ten percent autologous serum supplementation was considered to be the closest substitution to physiological conditions for moDC cultivation. Research teams studied moDC yield, maturation, antigen presentation, and ability to prime lymphocytes, cytokine production and many other functions when autologous serum was used. Most of these studies concluded that serum free medium is the most suitable for clinical trials^2–7^. Some of the papers used also autologous plasma in their experiments. However, not all of them mention how the plasma was obtained or what was its concentration.

Thereafter we tested the effect of autologous serum and plasma from EDTA coated tubes on peripheral blood mononuclear cells (PBMC) cultivation. Our aim was to determine the effect of autologous serum and plasma preferably on monocytes as a source for moDC generation. We found out that after 24 hours there was a decrease in number of monocytes when cultivated in RPMI supplemented with plasma from EDTA coated tubes as compared to PBMCs cultivated in RPMI supplemented with autologous serum or FBS. After 48 hours the decrease was even more profound, monocytes virtually disappeared from culture when EDTA plasma was used (Figure 2A). We observed similar results in lymphocyte population. Additionally, proliferation of CD4+ and CD8+ lymphocytes was also affected, with impaired proliferation in media supplemented with autologous plasma obtained from EDTA coated tubes. The lymphocytes did not restore their capacity for proliferation when ions, Mg^2+^ and Ca^2+^were added to the plasma.

**Figure 2:**
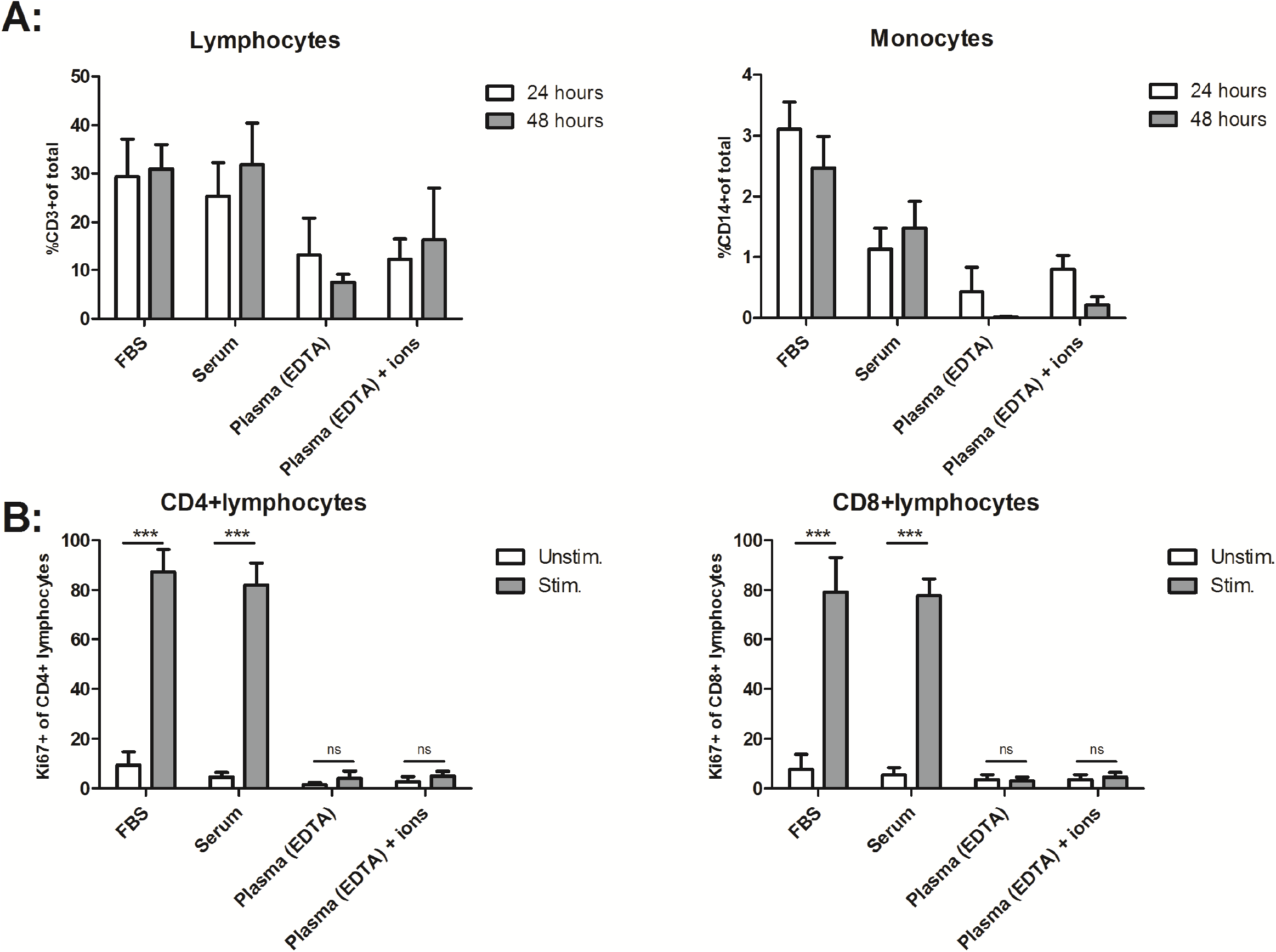
(A) Peripheral blood mononuclear cells (PBMCs) were cultivated in RPMI supplemented with FBS, autologous serum or plasma and proportion of CD3+ and CD14+ cells were analyzed after 24 and 48 hours. (B) Additionally, proliferation of CD4+ and CD8+ lymphocytes were tested after 72 hours after stimulation with antiCD3-antiCD28. The assay was performed in triplicates and mean values +/− SEM are shown.

The effect of autologous plasma on lymphocyte proliferation was studied many years ago. Plasma used in these early experiments was mostly obtained from heparinized blood^12–14^. Nevertheless, heparin is a biologically active molecule which modulates many immunological processes^15^. Therefore, we tried to avoid using plasma from heparin coated tubes in our experiments, as the stimulatory capacity, cytokine production and other functions of blood cells depend on which anticoagulant the blood was drawn into. EDTA seems to decrease the functional capacity of these cells in whole blood assays^16^.

We have found out that autologous plasma obtained from EDTA coated tubes is absolutely unsuitable for cell culture media. When this plasma was present in culture media, viability of moDC and PBMC was decreased and lymphocytes were unable to proliferate. Our findings suggest that myeloid cells are more sensitive to supplementation used in culture media and the best chance to provide successful experiments is to avoid using plasma from EDTA coated blood tubes. We strongly recommend using autologous serum instead of plasma in cell cultures with autologous conditions.

## Acknowledgement

The study was supported by the Czech Ministry of Health AZV 16-32838A.

